# Gene expression signatures predict response to therapy with growth hormone

**DOI:** 10.1101/637892

**Authors:** Adam Stevens, Philip Murray, Chiara De Leonibus, Terence Garner, Ekaterina Koledova, Geoffrey Ambler, Jia-Woei Hou, Klaus Kapelari, Jean Pierre Salles, Gerhard Binder, Mohamad Maghnie, Stefano Zucchini, Elena Bashnina, Julia Skorodok, Diego Yeste, Alicia Belgorosky, Juan-Pedro Lopez Siguero, Regis Coutant, Eirik Vangsøy-Hansen, Lars Hagenäs, Jovanna Dahlgren, Cheri Deal, Pierre Chatelain, Peter Clayton

**Author notes:** These authors contributed equally to the work (Adam Stevens, Philip Murray).

## Abstract

Recombinant human growth hormone (r-hGH) is used as a therapeutic agent for disorders of growth including growth hormone deficiency (GHD) and Turner syndrome (TS). Treatment is costly and current methods to model response can only account for up to 60% of the variance. The aim of this work was to take a novel genomic approach to growth prediction. GHD (n=71) and TS patients (n=43) were recruited in a study on the long term response to r-hGH over five years of therapy. Pharmacogenomic analysis was performed using 1219 genetic markers and baseline blood transcriptome. Random forest was used to determine predictive value of transcriptomic data associated with growth response. No genetic marker passed the stringency criteria required for predictive value. However, we demonstrated that transcriptomic data can be used to predict growth with a high accuracy (AUC > 0.9) for short and long term therapeutic response in GHD and TS. Network models identified an identical core set of genes in both GHD and TS at each year of therapy whose expression can be used to classify therapeutic response to r-hGH. Combining transcriptomic markers with clinical phenotype was shown to significantly reduce predictive error. We have characterised the utility of baseline transcriptome for the prediction of growth response including the identification of a set of common genes in GHD and TS. This work could be translated into a single genomic test linked to a prediction algorithm to improve clinical management.

**One Sentence Summary:** A blood transcriptome signature predicts response to recombinant human growth hormone in both growth hormone deficient and Turner syndrome children

**Trial registration numbers:** NCT00256126 & NCT00699855

## Introduction

Recombinant human growth hormone (r-hGH) is used as a therapeutic agent for a range of disorders of growth impairment including growth hormone deficiency (GHD) and Turner syndrome (TS). Treatment is costly at between £6000 – £24000 per centimetre (cm) gained in final height (*1*). Therapy is not always successful in patients and there are currently no genomic markers for predicting positive or negative responses. Prediction models up to four years of therapy have been defined using clinical measurements (*2*) but have been difficult to implement in practise. Whilst an understanding of the pharmacogenetic background has been established (*3, 4*), such approaches are of limited predictive value due to the influence of covariates related to the child’s developmental stage, disease severity and geographical location (*5*, 6). The pre-treatment blood transcriptome has been previously shown to relate to first year response to r-hGH therapy (*7*), however, little is known about the predictive value of this association and its relationship to longer term response to therapy. The transcriptome represents a level of ‘omic’ data that reflects genetic information, developmental stage in relation to age (*8*) along with the impact of the local environment (*6*) and, therefore, has potential to classify response to r-hGH.

Response to r-hGH in the first year of therapy is considered to be a primary marker of growth response. Prediction of first year growth has been shown to be dependent on GHD severity, age, distance to target height, body weight, dose of r-hGH, birth weight and, as defined by regression models, can account for 61% in GHD (*9*) and 46% in TS (*10, 11*) of the variation within the data. Clinical markers such as distance to target height are surrogate genetic variables and this implies that an effective level of genomic prediction is hypothesised to be possible if developmental (*8*, 12) and environmental covariates (*13*) of growth response can be taken into account.

Transcriptomic data have been used extensively in cancer tissues both to sub-type the tumour (*14-16*) and to predict response to therapies (*17*, 18). In contrast in this study we have used peripheral blood gene expression profiling as the source for gene expression profiles, and show that these patterns can be used to predict response to r-hGH in each year of treatment up to five years in two different growth disorders that account for approximately 60% of GH prescriptions.

## Results

### Growth response of patients over five years of r-hGH treatment

The auxology of the PREDICT study has been previously described at baseline and after one year (*7*) and after three years (*5*) of therapy with r-hGH. Height velocities as a measure of response to r-hGH at each year in GHD and TS are shown in **Table 1A**. As expected, first year growth response is the largest with a decline in subsequent years to a maintenance growth rate (*19*).

**Table 1.** Patient characteristics. **A)** Growth response endpoints used over the duration of the study and **B)** baseline auxology for patients with growth hormone deficiency (GHD) and Turner Syndrome treated with recombinant human growth hormone (r-hGH).

| Condition | Height Velocity<br>at year of treatment | Mean<br>(± standard deviation) | Median<br>(min, max) | N |
| --- | --- | --- | --- | --- |
| **GHD** | **HV1** | 8.9(±2.1) | 8.7 (4.7, 14.3) | 71 |
| | **HV2** | 7.4 (±1.6) | 7.1 (3.4, 12.2) | 65 |
| | **HV3** | 6.6 (±2.0) | 6.5 (2.0, 11.4) | 65 |
| | **HV4** | 6.1 (±2.3) | 6.2 (0.9, 11.6) | 60 |
| | **HV5** | 5.1 (±2.3) | 5.2 (0.0, 10.8) | 53 |
| **TS** | **HV1** | 7.6 (±1.4) | 7.2 (5.3, 11.7) | 43 |
| | **HV2** | 6.0 (±1.1) | 6.1 (3.3, 8.0) | 31 |
| | **HV3** | 5.3 (±1.5) | 5.0 (1.9, 8.2) | 40 |
| | **HV4** | 4.7 (±1.8) | 4.8 (1.1, 8.1) | 41 |
| | **HV5** | 3.7 (±1.6) | 3.9 (1.0, 7.4) | 33 |

| Clinical Characteristics | GHD (N=70) | TS (N=43) |
| --- | --- | --- |
| Male | 45 (64.3)* | 0 (0.0)* |
| Female | 25 (35.7)* | 43 (100)* |
| Age at baseline (years) | 9.3 (6.0, 11.2) | 9.9 (7.2, 11.8) |
| Baseline height SDS | -2.1 (-2.5, -1.7) | -2.5 (-3.2, -1.9) |
| Baseline BMI SDS | -0.2 (-0.9, 0.3) | 0.4 (-0.3, 1.2) |
| MPH SDS | -0.7 (-1.5, 0.0) | -0.1 (-0.9, 0.6) |
| GH peak response (μg/L) | 3.9 (2.3, 5.6) | - |

Data are n (%) or median (Quartile 1, Quartile 3). \*All were Tanner Stage 1 at baseline. BMI, body mass index; GH, growth hormone; GHD, growth hormone deficiency; TS, Turner syndrome; MPH, mid-parental height; SDS, standard deviation score.

### Genetic associations were not robust enough to be used to predict changes in growth rate over the five years of the study

The association between SNP carriage and growth response was assessed for 1096 and 792 growth-related candidate genes, in GHD and TS respectively, which passed the filtering criterion. Whilst 113 SNPs were associated with growth response endpoints with an FDR p-value <0.05 modified by the number of blocks of linkage disequilibrium, none of these were deemed to pass the stringency criteria required for predictive value (**Supplemental Table S1A-E**).

### Unsupervised and supervised analysis demonstrates that GHD and TS blood transcriptome at baseline can be used to classify response to r-hGH therapy over five years of treatment

We first demonstrated that a fundamental relationship existed between the baseline blood transcriptome and response to r-hGH over the 5 years of the study (GHD n= 50, TS n=22) using DAPC on the unsupervised baseline transcriptome (GHD = 8875, TS = 8455 gene probe sets). These analyses showed clear segregation of the low response (LoQ) and the high response (HiQ) quartiles of response to r-hGH thus demonstrating the utility of blood transcriptome to differentiate response groups (**Figure 1**). Partial least squares Discriminant Analysis (PLS-DA) of the unsupervised baseline transcriptome demonstrated similar findings (**Figure 2**).

**Figure 1.**
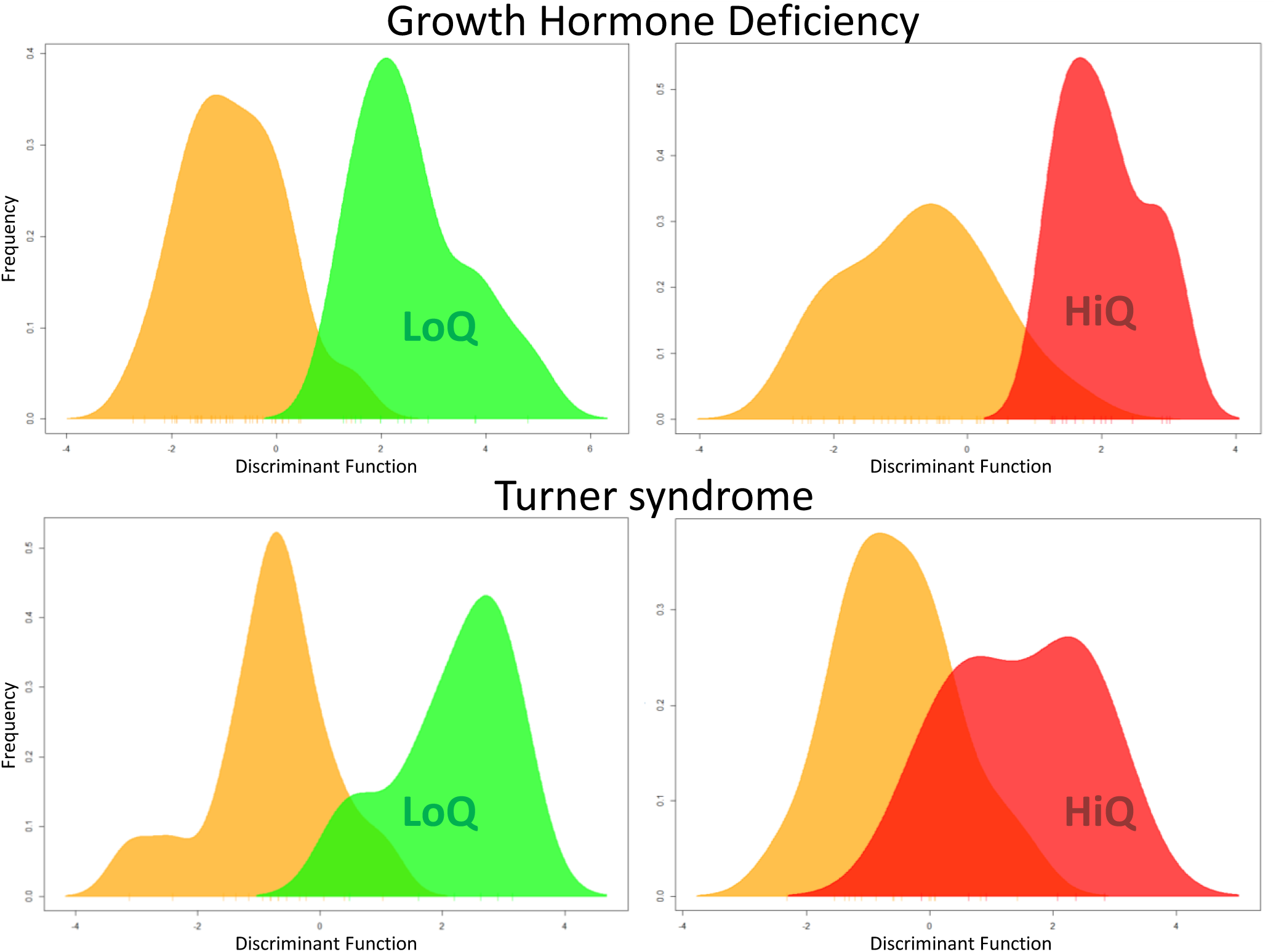
The association of whole blood gene expression at baseline with response to recombinant human growth hormone (r-hGH) over all five years of therapy in patients with growth hormone deficiency (GHD) and Turner syndrome (TS). Comparison of patient response to r-hGH using Discriminant Analysis of Principal Components (DAPC). Low quartile (green, LoQ) and high quartile (red, HiQ) of growth response over five years of therapy (cms grown) compared to the remaining patients (orange) in GHD (N= 50) and TS (N=22). Unsupervised transcriptomic data with no normalisation for phenotype are shown, GHD = 8875 & TS = 8455 gene probesets. DAPC generates a discriminant function, a synthetic variable that optimises the variation between the groups whilst minimising the variation within a group. The frequency of the discriminant function of DAPC is plotted.

**Figure 2.**
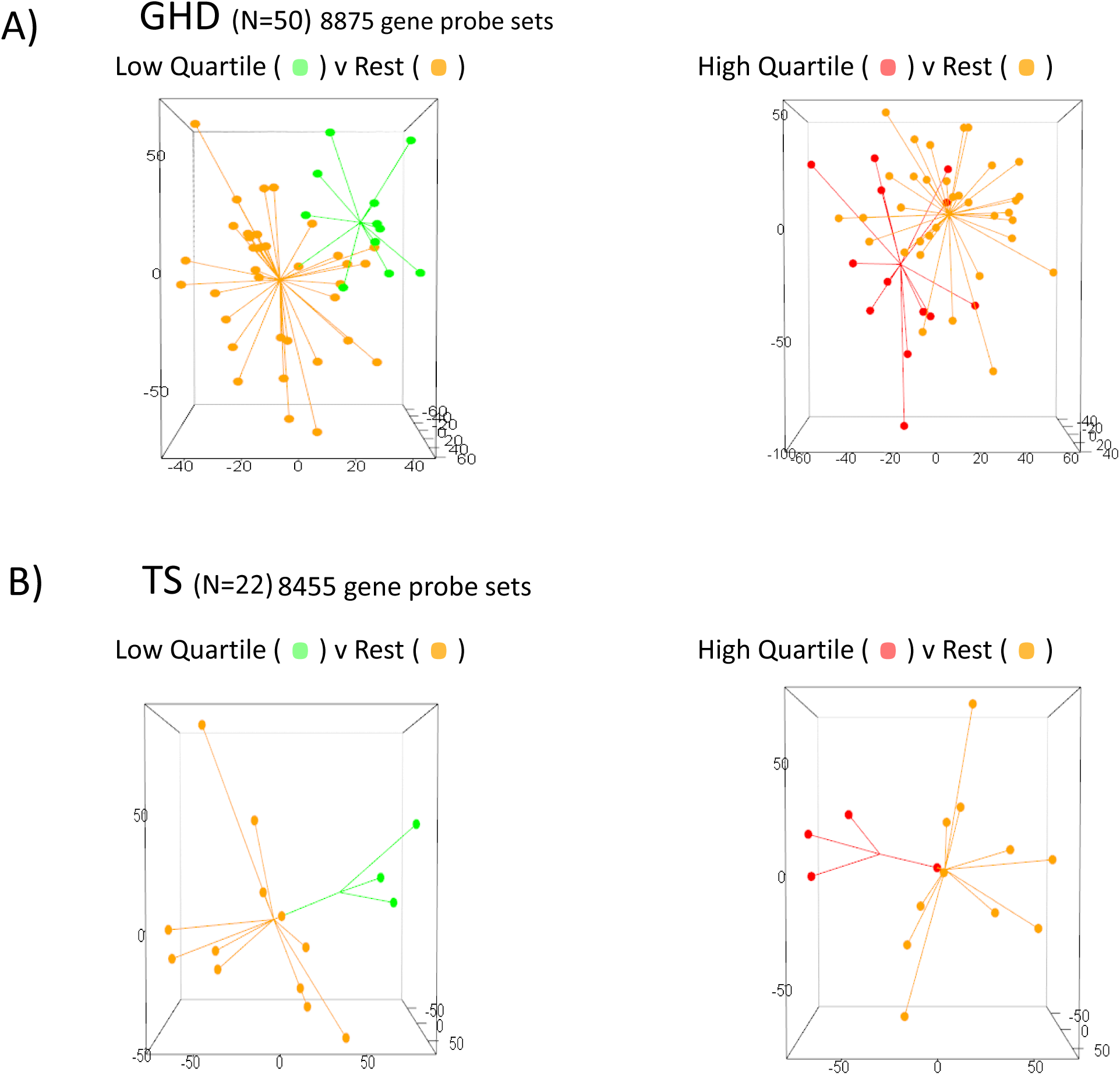
Whole blood gene expression is associated with response to recombinant human growth hormone (r-hGH) over five years of therapy in patients with growth hormone deficiency (GHD) and Turner syndrome (TS). Partial least squares discriminant analysis (PLS-DA) of unsupervised transcriptome using three components. The low and high quartiles of growth response are shown for response to r-hGH (cm) over five years in **A)** GHD and **B)** TS. Star plot shows sample distance from the centroid, the arithmetic mean position of all the points in each group.

### GHD and TS blood transcriptome at baseline can be used to classify response to r-hGH therapy year-on-year over five years of treatment

Baseline gene expression associated with height velocity at each year of the five years after the start of treatment with r-hGH was defined using rank regression (p<0.01) **(Supplemental Table S2)** with a range of covariates – microarray batch, age, body mass index (BMI) at baseline for both GHD and TS patients along with gender and peak GH test response in GHD. Tanner stage was added as a further covariate to account for the pubertal status of the patients **(Figure 3).** There was no difference in auxology at baseline between each group of patients at each year of the study **(Table 1B & Supplemental Table S2).** First classification of low and high responding quartile groups of patients was assessed by PLS-DA using unmodified class sizes (**Supplemental Table S3A & 3B**): clear separation of the quartiles was observed (example of first year GHD response, **Figure 4**). We also examined classification of growth response using random forest (RF) with oversampling by SMOTE to correct for uneven class size (GHD, **Supplemental Table S3A** and TS, **Supplemental Table S3B**). These data show clear classification of good and poor responders: at each year of the study all PLS-DA area under the curve of the receiver operating characteristics (AUCs) were between 73% and 98% and all RF AUCs were between 78% and 98% in both conditions.

**Figure 3.**
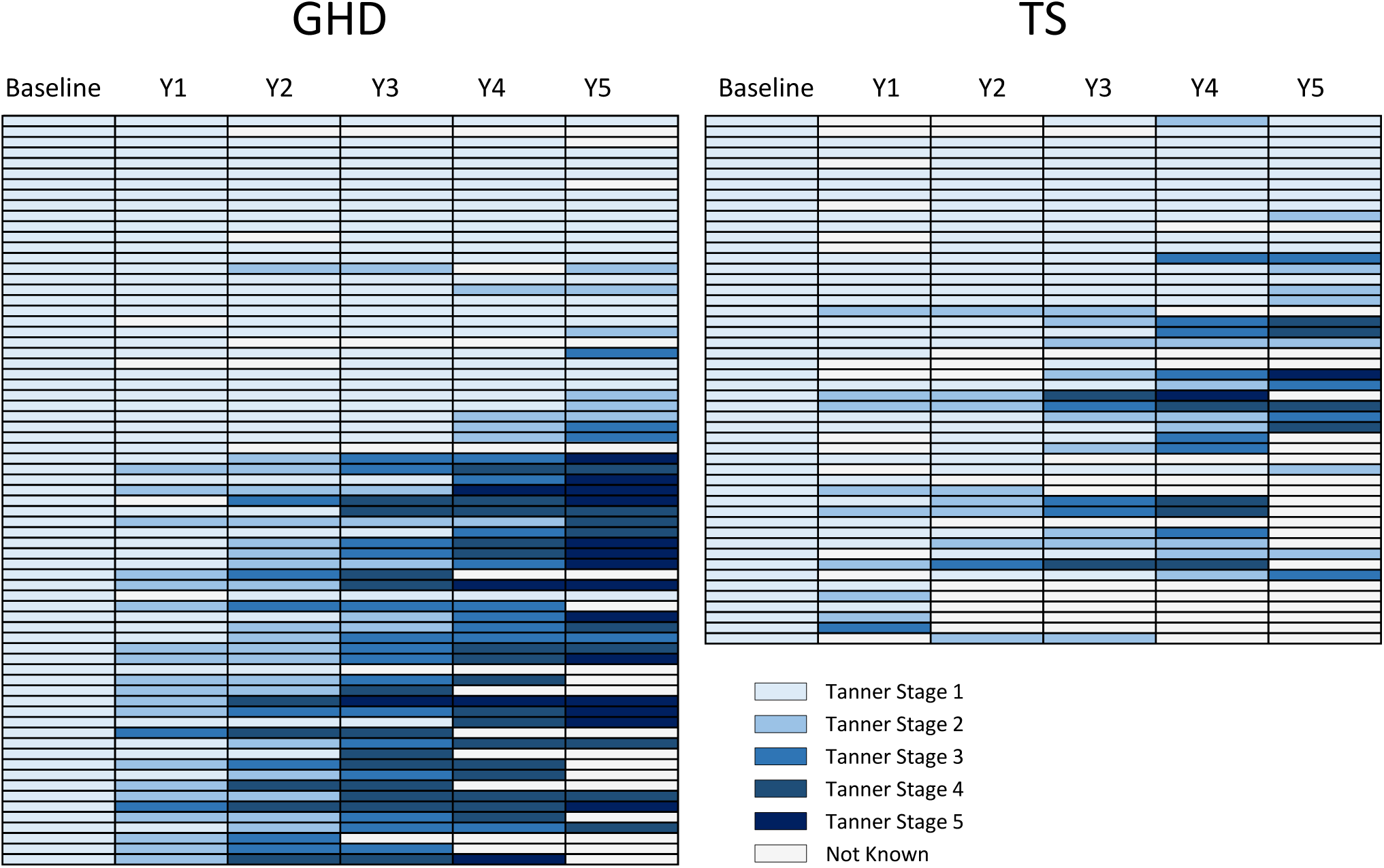
Distribution of Tanner stages over the study duration. Heat map of the Tanner stage of each patient (row) ordered by age (youngest at top). Y = year of study.

**Figure 4.**
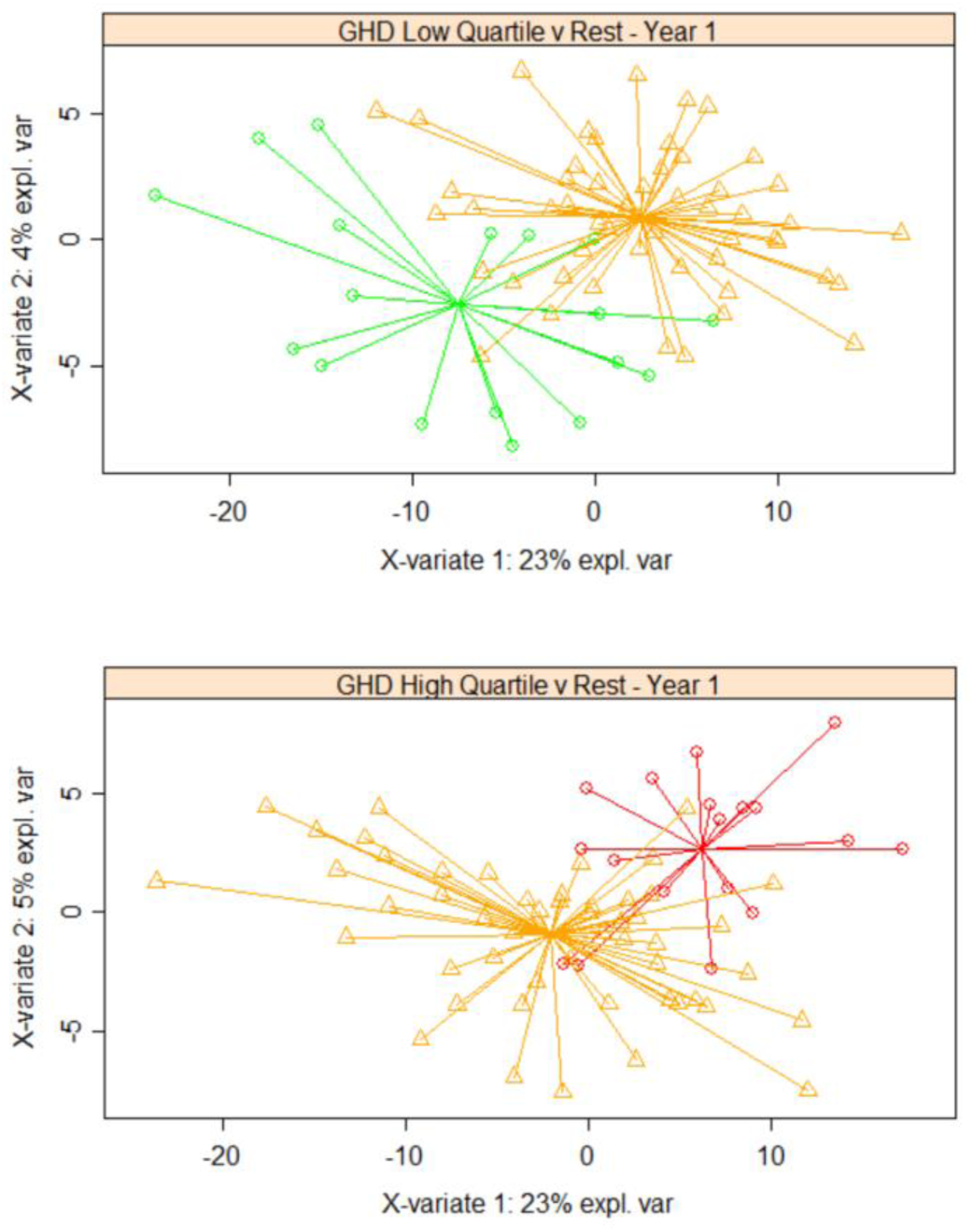
Predictive value of whole blood gene expression associated with response to recombinant human growth hormone (r-hGH) in patients with growth hormone deficiency (GHD). Classification of low quartile (LoQ) and high quartile (HiQ) of growth response (height velocity, cm/year) over each of five years of therapy with r-hGH (Y1-Y5) was performed in GHD patients and TS patients. Gene expression associated with growth response was determined using rank regression (p<0.01) and Partial least squares discriminant analysis (PLS-DA) with two components (X-variate 1 & 2) was used to visualise response groups; PLS-DA is an analytical approach that determines the similarity between individual patients whilst maximising the difference between patient groups. Low quartile (green) and high quartile (red) compared to the rest of the data (orange) is shown for first year growth response to r-hGH in GHD (N = 71, 330 gene probesets with rank regression p<0.01). Similarity between samples is represented by their proximity. The star plot shows sample distance from the centroid, the arithmetic mean position of all the points in each group.

### Interactome network models of response to r-hGH

There was limited overlap between GHD and TS whole blood transcriptomic markers related to growth response at each year of the study (**Supplemental Table S2**). We therefore generated interactome network models including inferred interactions to assess whether GHD and TS growth response-associated gene expression was related by affecting the function of similar network modules, albeit in different ways.

Interactome network models of gene expression associated with height velocity at each year of the study were generated. The hierarchy of overlapping modules of genes was identified in each network using the network topology parameter of “centrality” **(Supplemental Table S4)**. Network centrality is measurement that is known to be related to gene function within networks; the more central a gene is, the more capable it is of influencing other genes within the network (*20*).

The gene level summary of SNP associations with change in height and height velocity measurements with FDR <0.05 (**Supplemental Table S1C and S1D**) were mapped onto the network models **(Supplemental Table S4)**. Most of the genetic associations with change in height and height velocity were also present within the network models – 15/25 SNPs in GHD and 9/12 in TS (**Supplemental Table S5**), implying that these genes have a functional role in network action.

Network models associated with height velocity in each year in both GHD and TS demonstrated significant overlaps (Hypergeometric test, p<0.01) (**Figure 5)**. These observations imply that whilst associated gene expression may be different between GHD and TS, common network elements are being affected in the two conditions.

**Figure 5.**
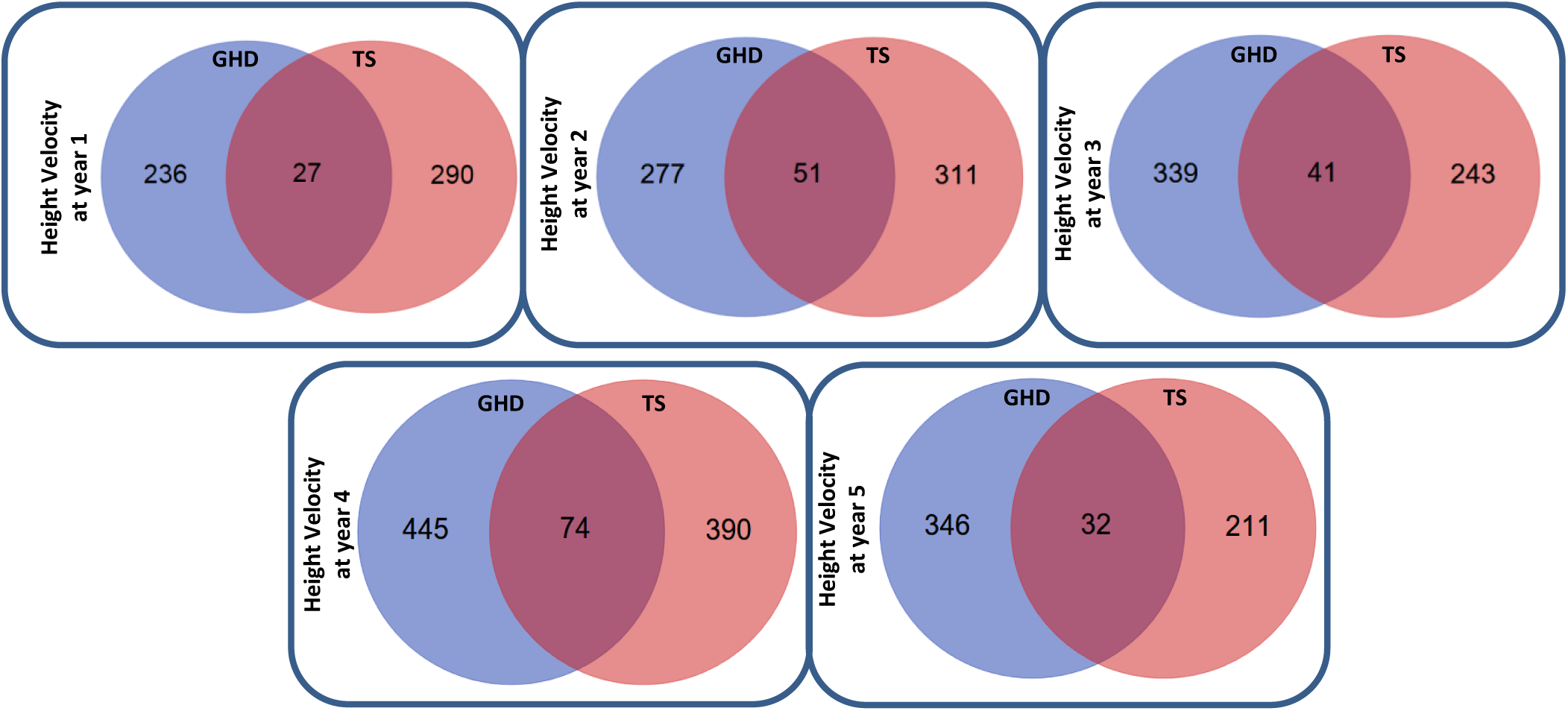
Overlap of the core interactome models of height velocity related gene expression in GHD and TS. Interactome models were generated from the gene expression associated (p<0.01) with the height velocity at each year of the study. The functional hierarchy of gene interaction modules within the interactome models was determined using the Moduland algorithm and the core of the interactome model was defined as the unique sum of the top ten elements of the modules as ranked by network centrality. The overlap of the core of the interactome models between GHD and TS was then determined and visualised as a Venn diagram.

The overlap between network models formed a discrete interactome element shared between GHD and TS (**Figure 6**). When this network was partitioned into genes related to each year of response to r-hGH (coloured **Figure 6A**), it was determined that the genes associated with year 3 formed a less distinct cluster within the network (**Figure 6B**). This observation is in alignment with a partition between early (years 1 and 2) and later (years 4 and 5) response to r-hGH as would be expected clinically.

**Figure 6.**
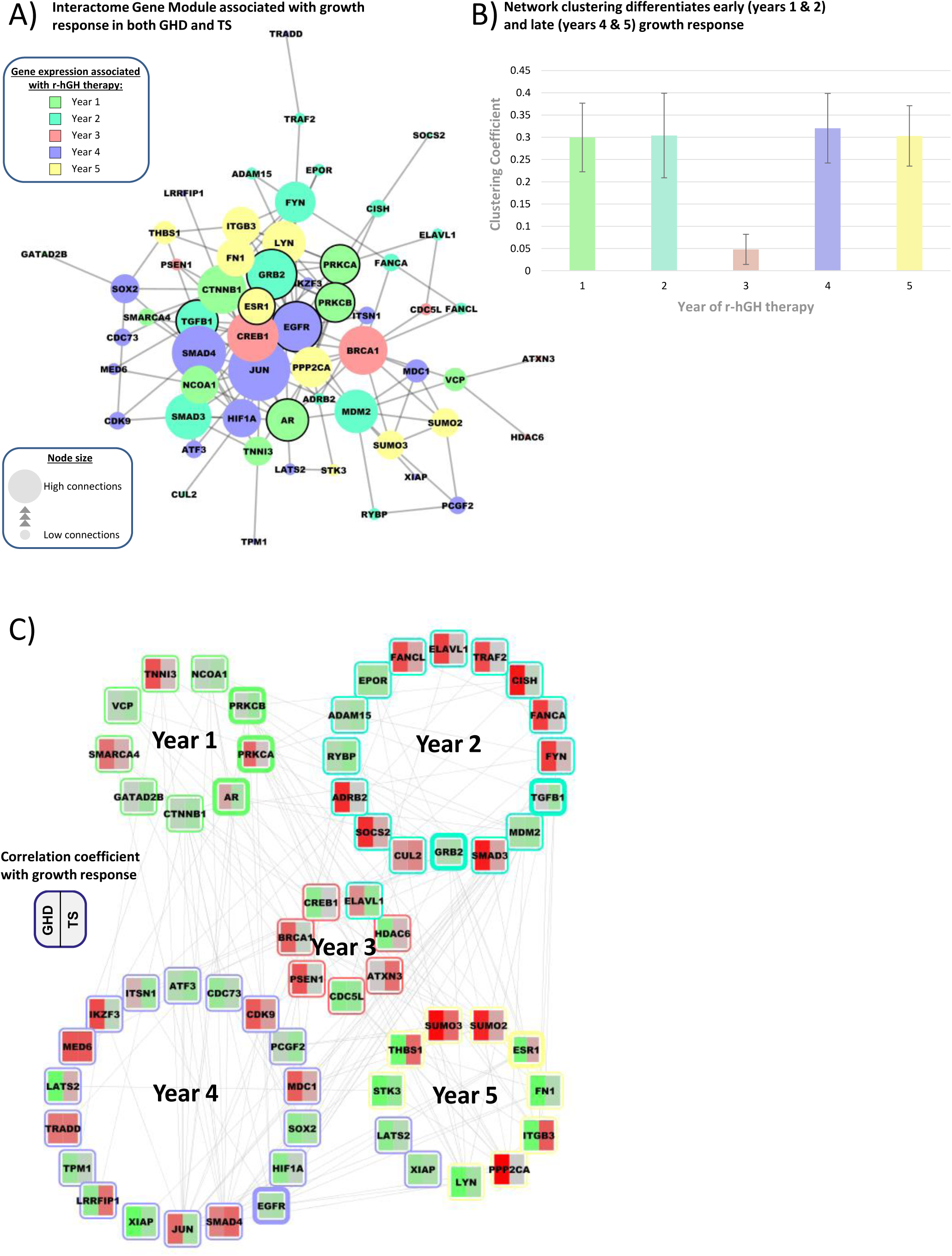
Network structure of the common core network module shared in patients with growth hormone deficiency (GHD) and Turner syndrome (TS) related to response to recombinant human growth hormone (r-hGH). **A)** Similarities in the interactome models of the response of GHD and TS to r-hGH were identified by overlap at each year of therapy. Genes were selected that were significantly related to growth response in either or both GHD and TS. The genes related to each year of therapy were combined into a set of 58 uniquely identified genes and this set was used to generate an interactome module (Reactome plugin for Cytoscape 3.6.0). Genes with a dark border also have a genetic association with growth response in either GHD or TS. Connecting lines represent known protein:protein interactions, size of the node is proportional to the number of connections made. **B)** The clustering coefficient of the group of genes in the network module associated with each year of therapy was determined and presented as a histogram (average ± standard error of the mean). The clustering coefficient measures the tendency of nodes to cluster together within a network. **C)** The correlation coefficient linking gene expression with growth response at each year of therapy was mapped to the network model, red = positive correlation, green = negative correlation. Genes with a thick border also have a genetic association with growth response in either GHD or TS.

The facts that **i)** genetic associations with growth response map to the network models derived from transcriptomic data and that **ii)** the network connectivity of the central modules changes over the duration of the study imply that the network models are robust and account for the effect of development on related phenotype (**Supplemental Table S5**).

### The Identification of core sets of genes that can classify response to r-hGH in both GHD and TS

The overlap between network models was used to select a common set of genes at each year of therapy present in both GHD and TS. Genes within this common list were selected for growth response classification if they had previously been identified as significantly associated with height velocity by rank regression in either GHD or TS (p<0.05) (**Figure 6C** & **Supplemental Table S2**).

Classification of both high and low r-hGH response quartiles against the remaining patients was shown using PLS-DA (no oversampling) and RF (using SMOTE oversampling). All AUCs for classification were between 74% and 96% (**Supplemental Table S6**).

Further confidence in the findings was provided by assessing the predictive quality of the gene probe sets using BORUTA to define the limits of the noise in the analysis using a 100-fold permutation of the data (e.g. first year growth response **Figure 7** & **Supplemental Table S7**).

**Figure 7.**
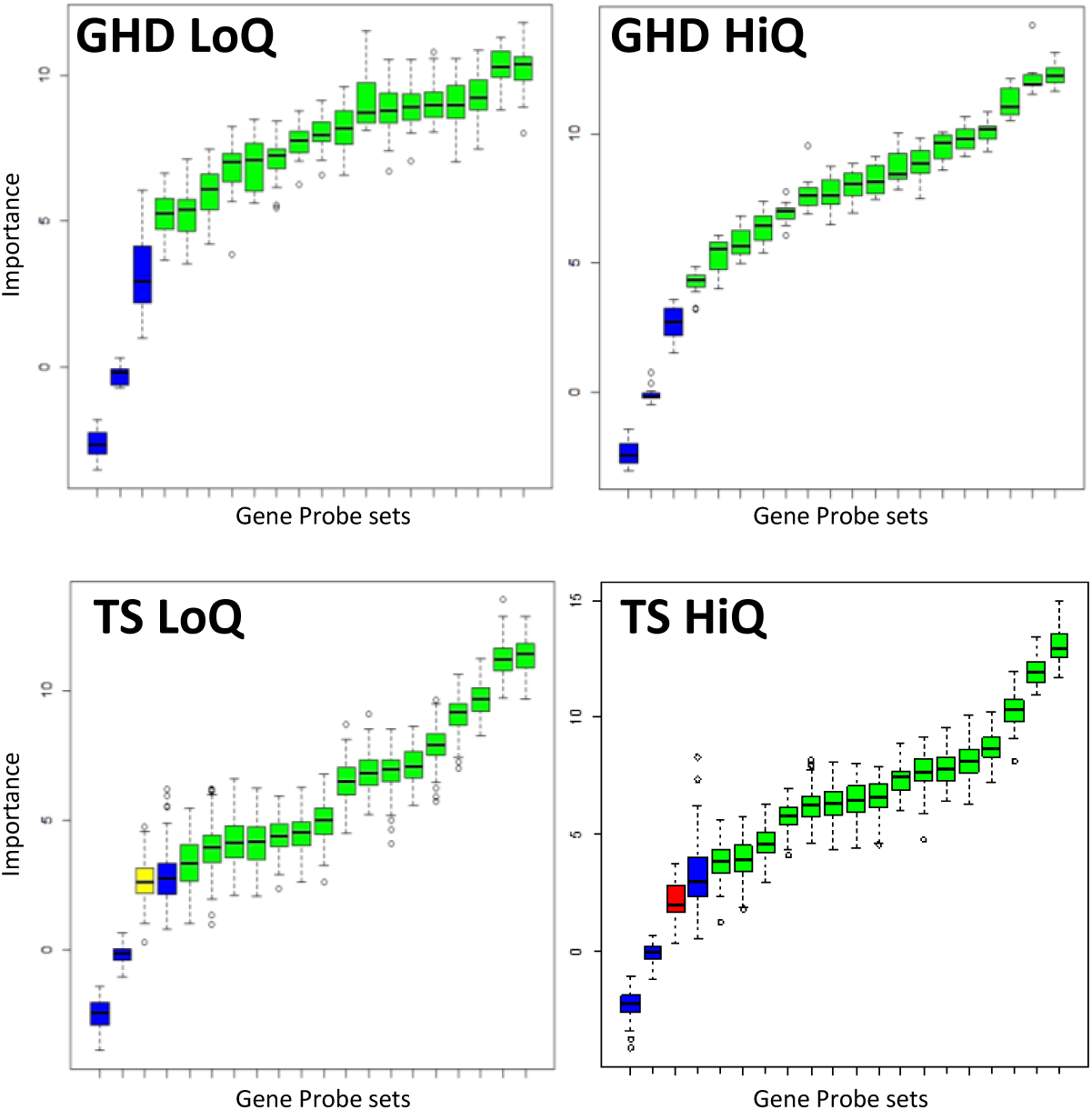
Predictive value of an identical set of blood gene expression markers identified by network analysis in the classification of response to recombinant human growth hormone (r-hGH) in patients with growth hormone deficiency (GHD) and Turner syndrome (TS). First year growth response is used as an example. Similarities in the interactome models of the response of GHD and TS to r-hGH were identified by overlap at each year of therapy. Genes were selected that were significantly related to growth response in either or both GHD and TS, generating an identical set of gene probesets used for prediction of both high and low response in both GHD and TS. BORUTA, an all relevant feature selection wrapper random forest based algorithm, was used to confirm the importance of gene expression probe-sets used for classification of response to r-hGH. The BORUTA algorithm uses a 100-fold permutation to define the noise present in the data; the noise is modelled as shadow variables and used as a basis to assess confidence in the data. Green = confirmed gene probeset, yellow = tentative gene probeset, red = rejected gene probeset, blue = shadow variables (high, medium and low shadow variables are derived to define the noise within the dataset). Low quartile (left column-LoQ) and high quartile (right column-HiQ) are shown for first year growth response to r-hGH in GHD and TS. The same group of gene probesets are used in each case.

### The core sets of genes with expression in whole blood that can classify response to r-hGH in both GHD and TS are associated with differential genomic methylation

Changes in genomic methylation in response to short term treatment with r-hGH (4 days) have been demonstrated in children with range of conditions that manifest short stature (*21*). Using the data provided by this previously published study we examined the epigenome at baseline (prior to r-hGH treatment) in relation to growth response (measured by knemometry) in GHD patients (n=6) and found that using a gene level summary of DNA methylation (20618 genes) 497 had methylation associated with growth response to r-hGH (rank regression, p<0.01) (**Figure 8**). The majority of associated genes (425/497) were hypermethylated at lower rates of growth response.

**Figure 8.**
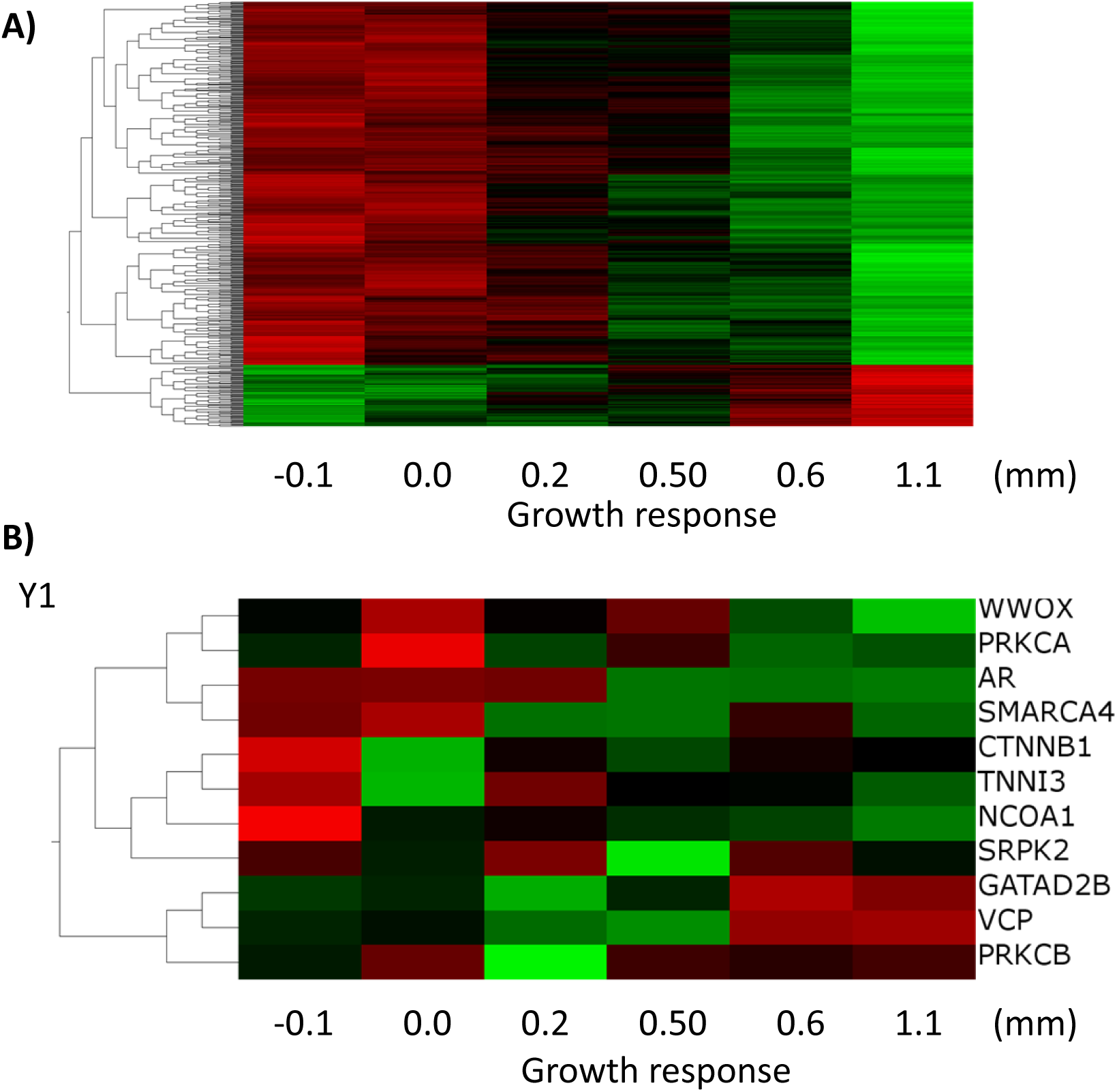
Gene level summary of DNA methylation in GHD patients is related to growth response as measured by Knemometry. Whole epigenome measurements of six GHD patients with growth response after 4 days of r-hGH therapy measured by knemometry were available from previously published data (GSE57107). A gene level summary of DNA methylation was conducted using median values in Qlucore Omics Explorer (version 3.3) (n=20618). **A)** Rank regression of whole genome DNA methylation against growth response after 4 days of r-hGH as measured by knemometry (p<0.01) found 497 genes with differential methylation the majority of which showed increased methylation at low rates of growth (negative correlation). **B)** Whole genome methylation in the six GHD patients ordered by growth response in the sets of genes identified as predicting response to r-hGH in the first year of therapy in both GHD and TS.

We took the core sets of genes previously identified as classifiers of response to r-hGH in both GHD and TS and mapped gene level methylation present in the six GHD patients with knemometry measurements. The majority of genes (57/71) were correlated with growth response (|R|>0.3) these were evenly distributed between positive (n=27/71) and negative (n=30/71) correlations (year one data shown in **Figure 8B**).

### Transcriptomic markers combined with phenotype lead to better growth response prediction

It is known that the baseline phenotype of GHD and TS patients can be used to predict response to r-hGH (*2*, 22-24). We found that including the blood transcriptome markers increased predictive value at each year by an average of 7% (p=0.0031) and 4% (p=0.0365) (prediction of low quartile) along with 4% (p=0.0179) and 4% (p=0.0097) (prediction of high quartile) in GHD and TS respectively (**Table S8**).

Importantly we also noted a significant decrease of error rate in the prediction of growth response at each year when blood transcriptome markers were combined with clinical phenotype markers. Error rates decreased by an average of 5% (p=0.0084) and 5% (p=0.0400) (prediction of low quartile) along with 5% (p=0.0252) and 5% (p=0.0067) (prediction of high quartile) in GHD and TS respectively (**Table S8**). The reduction observed amounted to an average halving of the error rate seen when predicting response to r-hGH using clinical phenotype markers alone.

## Discussion

This study aimed to identify for the first time the genomic associations that classify response to r-hGH therapy from one year up to five years of treatment with r-hGH in children with TS and GHD.

Our previous analysis has shown limited utility of genetic associations derived from a candidate set of growth related genes in the prediction of response to r-hGh in GHD and TS after one year of therapy (*5*, 7, 25). Hence genetic data do not appear to be powerful enough on their own to be used in prediction and clinical management.

The whole blood transcriptomic profile of GHD and TS patients has been shown to be associated with first year growth response to r-hGH (*7*) and to correlate with the interaction between GHRd3 and GHD severity (*25*). We therefore reasoned that there may be value in using transcriptomic data to classify growth response, as it reflects both a child’s genetic profile and the complex clinical phenotypes arising from changes in physical development during childhood, as well as variation in the severity of the underlying condition. By normalising gene expression for phenotype, including pubertal stage, we were able to show that whole blood transcriptomic data, associated with height velocity at each year of the study, could be used to classify both the low and high quartiles of growth response, with ‘Area under the Curve’ up to 97%, providing the basis for a predictive test.

Little overlap between GHD and TS was observed between the gene expression data that was associated with each year of growth response. We therefore investigated whether GHD and TS were interacting with similar functional units of genes using network models (*26*). We generated network models of growth response (as determined by height velocity) at each of the five years of treatment using baseline gene expression. Functional modules of genes within these models were ranked according to their network centrality. The measure of network centrality is known to be associated with mechanism (*27, 28*) and we used this measure to define the functional hierarchy of the modules of genes whose expression was linked to r-hGH response at each year of therapy.

We demonstrated robustness of the network modules identified by mapping the genetic associations identified in this study to the network models. This process highlighted genes previously identified as associated with growth response after one year of therapy (*GRB10, SOS1* and *INPPL1* in GHD) (*7*) along with one month change in serum IGF-I associated with r-hGH therapy (CDK4) (*29*). It was also noted that three genes were present (*INPPL1* and *SOS1* in GHD and *PTPN1* in TS) out of the four genes identified within the PREDICT validation study as having replicated an association with first year growth response when controlled for co-variates (*5*).

A significant overlap between the core network gene modules between GHD and TS was identified. We then used gene expression changes associated with growth response within these network elements to identify genes common to both conditions and show that their expression could be used to classify growth response. The major strength of this study is to have identified predictive markers and common genomic mechanisms related to early and later growth in two different growth disorders. Our findings are also supported by the demonstration of differential methylation in these shared genes, associated with response to r-hGH in another study (*30*). Importantly we have defined sets of gene expression with predictive value in two conditions where the number of genes (17–26) is smaller than the number of patients in the group (33-70) [**Table S2** & **S6**]; this indicates that the findings are not a consequence of overfitting (*31*).

In this study we have compared the use of baseline patient auxology to blood transcriptome in predicting response to r-hGH. Linear models based on baseline patient auxology can account for ∼40-60% of the variance observed (*9, 10*). Using random forest we found no significant difference in the AUC of baseline auxology alone compared to using blood transcriptome alone in either GHD or TS (all ∼90%). It should be noted that this comparison was with the transcriptome shared between GHD and TS and if the full blood transcriptome is used then the average AUC is significantly higher than that derived from baseline auxology (average AUC ∼90% compared to ∼95%). We recognise that further work would need to be done to refine a smaller number of genes and therefore minimise the risk of overfitting when using the full blood transcriptome. However, we did identify a significant boost to prediction of between 4% and 7% when the transcriptomic signature shared between GHD and TS was combined with the baseline patient auxology. Importantly the gain in prediction was combined with an average halving of the error rate, a feature that represents a major clinical advance in the prediction of response to r-hGH.

This work has led to three novel findings relevant to growth studies, and potentially to other therapeutic areas in paediatrics. First, this study has demonstrated the utility of whole blood transcriptome in the classification of growth response in GHD and TS, derived from a baseline blood sample which is straightforward to obtain in any child. This technique may be of particular use in conditions with marked variability in response to r-hGH such as the short child born small for gestational age. Second, network analysis provides a novel approach that can be used to identify genomic features that are likely to have high predictive value. Finally, a set of common genes in GHD and TS identified by a network approach can be used to classify growth response in both conditions, providing the opportunity to develop a test to inform clinical management.

## Methods

### Patients

The PREDICT Long Term Follow up study (multicentre, open-label, prospective, phase IV) and the pharmacogenetics of the first year of r-hGH treatment have been described extensively previously (*7*, 29). Briefly, pre-pubertal children with GHD and TS were enrolled. A diagnosis of GHD was reached following two pharmacological stimulation tests with a peak GH concentration of < 10µg/L. Prior to enrolment in the study none of the children had received GH therapy. Children with GHD due to central nervous system tumours or radiotherapy were excluded but children born small for gestational age were not. The diagnosis of TS was based on karyotype.

This PREDICT study was conducted in compliance with ethical principles based on the Declaration of Helsinki, the International Conference on Harmonization Tripartite Guideline for Good Clinical Practice, and all applicable regulatory requirements.

### Genetic Analysis

A total of 1219 genetic markers were used in the analysis, 1217 Illumina-genotyped single nucleotide polymorphisms (SNPs) corresponding to a candidate list of 103 genes and 2 TaqMan-genotyped SNPs in the *IGFBP3* promoter. All genes selected are known to be involved in growth regulation and GH action as previously described (*5, 7*).

A Kruskal-Wallis rank sum test was applied on the following 3 genetic models **a)** genotypic (AA, AB, BB); **b)** dominant (AA/AB+BB) and; **c)** recessive (AA+AB/BB). For non-pseudoautosomal X chromosome markers, GHD boys and TS girls were analysed as having only two homozygote categories (AA/BB). Adjustment for multiple testing was performed using Bonferroni correction with 2 different parameters as the number of independent tests, the number of Linkage Disequilibrium (LD) blocks in the gene in which the SNP is contained and the total number of LD blocks present in all genes (768 in GHD and 563 in TS). Filtering criterion for prediction were defined as a false discovery rate [FDR] modified p-value <0.05 unmodified for linkage disequilibrium blocks.

### Transcriptome Analysis

Transcriptomic profiling was carried out on whole blood RNA as described previously (*7*) using Affymetrix GeneChip Human Genome U133 plus 2.0 Arrays. For background correction, the Robust Multichip Average (RMA) was applied with quantile normalisation and a mean probe set summarisation using Qlucore Omics Explorer 2.3 (Qlucore, Lund, Sweden). The data set generated was subject to quality control to investigate the presence of outliers and further confounding effects.

Baseline gene expression associations with height velocity in each year of growth response were determined using rank regression with microarray batch, age, body mass index (BMI) at baseline as covariates for both GHD and TS patients along with gender and peak GH test response (average of two provocative tests) for the GHD patients. Over the study a number of children either entered puberty spontaneously or received exogenous sex steroids for pubertal induction. We therefore introduced a further normalisation for Tanner stage to the analysis to account for the proportion of children entering puberty in each year of the study.

### Generation of network models

Network analysis allows the identification and prioritisation of key functional elements within interactome models. To derive an interactome model differentially expressed genes were used as “seeds” and all known protein:protein interactions between the seeds and their inferred immediate neighbours were calculated to generate a biological network using the output of the Biogrid model of the human Interactome (3.3.122)(*32*). Network generation and processing was performed using Cytoscape 2.8.3(*33*).

### Analysis of Gene Network Models

Clustering and “community structure” of modules within biological networks arise from variation in connectivity within the network and are known to be associated with function (*27*, 34). To rank these functional components within interactome models we used the ModuLand plugin for Cytoscape 2.8.3 to determine overlapping modules and to identify hierarchical structure using the centrality property thus enabling the identification of key network elements (*35*). The central core unit of each network module (metanode) was defined as the ten most central genes. A list of the unique genes in each metanode was generated and used as a model of the functional core of the associated network for further comparison. Network topology was analysed using the CytoHubba plugin for Cytoscape (*36*). The String database was used to assess the integrity and connectivity of gene modules (*37*).

### Analysis of epigenomic data

Epigenomic data from the whole genome DNA previously published methylation profiles of six GHD patients was used to assess the relationship of changes in DNA methylation in relation to response to r-hGH (*21*). The data from GSE57107 was re-analysed in Qlucore Omics Explorer 3.3 and a median based gene level summary of methylation was determined (n=20618). The relationship between gene level DNA methylation and response to r-hGH was determined using rank regression.

### Classification of Growth Response

All analysis was performed using the statistical software R 3.3.2 (*38*). The relationship of baseline gene expression to potential predictive value (classification of low and high quartiles of response) was performed using Discriminant Analysis of Principal Components (DAPC) (*39*), Partial Least Squares Discriminant Analysis (PLS-DA) (mixOmics 6.1.1 R package (*40*)) and random forest with 1000 trees (*41*). Class size imbalance was corrected for using Synthetic Minority Oversampling Technique (SMOTE) (*42*). Feature selection from random forest data was performed using the BORUTA algorithm (*43*). The area under the curve of the receiver operating characteristic (AUC) was used to present the probability of a randomly selected sample being classified correctly.

In random forests about one third of the cases are left out of each iteration and can be used as a test set to perform cross-validation and to get an unbiased estimate of the test set error, the out of bag (oob) error estimate. The oob error estimate is recognised as being unbiased (*41*).

We used random forest to investigate whether blood transcriptomic data from GHD and TS patients provided additional value for prediction of response to r-hGH based on baseline patient auxology (age, weight SDS, birthweight SDS and distance to target height SDS in both TS and GHD with the addition of peak GH value for GH provocation test in GHD). These analyses were performed by defining the predictive value of baseline clinical phenotype alone and these data were then compared to baseline clinical phenotype in addition to blood transcriptomic markers.

### Statistics

Analyses were performed to determine genetic associations with response to r-hGH using the Kruskal-Wallis rank-sum test with Bonferroni corrections for false discovery rate (FDR).

Transcriptomic data was subjected to dimensional scaling using Principal Components Analysis (PCA) and Iso-map multidimensional scaling (MDS) (*44*) and used to demonstrate data homogeneity (Qlucore Omics Explorer 3.3) along with outliers using cross-validation. Unsupervised analysis of transcriptome data was performed using a projection score to select optimal variable subsets by variance filtering (*45*).

Transcriptomic associations with response to r-hGH were performed using rank regression (p<0.01) and modified for the listed covariates. This was done by fitting a linear model with the factors to be eliminated as predictors, and retaining only the residuals (i.e. subtracting the part explained by the predictors). When a nominal factor was used as covariate (such as gender), this is equivalent to mean-centring each variable over each subgroup defined by the factor.

The significance of gene set overlaps derived from the network analysis was determined using the hypergeometric test. Analyses were performed in the stated software or using R (*38*).

### Study approval

The PREDICT (NCT00256126) and PREDICT long-term follow-up (NCT00699855) studies were approved by the Scotland Medical Research and Ethics Committee (reference 05/MRE10/61) and the North West Research Ethics Committee (reference 08/H1010/77), respectively. Informed consent was obtained from parents for all study participants.

## List of Supplementary Materials

**Supplement Tables.xlsx**

**Supplemental Table S1A.** Association of SNPs with growth response endpoints in GHD and TS.

**Supplemental Table S1B-D.** Gene level single nucleotide polymorphism (SNP) associations with growth response endpoints in growth hormone deficiency (GHD) and Turner syndrome (TS).

**Supplemental Table S2**. Transcriptomic associations with height velocity growth response endpoints at each year of the study.

**Supplemental Table S3.** Predictive value of whole blood gene expression associated with response to recombinant human growth hormone (r-hGH) in patients with growth hormone deficiency (GHD) and Turner syndrome (TS).

**Supplemental Table S4.** Interactome models of transcriptomic data associated with height velocity growth response endpoints at each year of the study.

**Supplemental Table S5.** Gene level SNP associations mapped to network properties of the interactome models of height velocity related gene expression.

**Supplemental Table S6.** Predictive value of an identical set of blood gene expression markers identified by network analysis in the classification of response to recombinant human growth hormone (r-hGH) in patients with growth hormone deficiency (GHD) and Turner syndrome (TS).

**Supplemental Table S7.** Overlap between GHD and TS interactome models at each year of therapy with r-hGH.

**Supplemental Table S8.** Comparison of the prediction of growth response using clinical phenotype with and without transcriptomic data.

## Acknowledgements

The following were PREDICT local centre principal investigators who enrolled patients into the study: Yang, S-W; Yoo, H-W; Wang, T-J; Cabrol, S; Weill, J; Pfaeffle, R; Antoniazzi, F; Pilotta, A; Veimo, D; Peterkova, V; Ferrandez Longas, A; Gonzalez, I; Quinteiro, S; Rodriguez Arnao, M; Bath, L; Colle, M; Brusquet, Y.

## Author Contributions

PEC and PC conceived and designed the PREDICT project and this study. Data analysis and methodology development were undertaken by AS, PGM, TG and CDL. The manuscript was written by AS and PGM and revised by EK, PC, and PEC. EK, GA, JH, KK, JPS, GB, MM, SZ, EB, JS, DY, AB, JPLS, RC, EVH, LH, JD, CD all contributed to the clinical data analysis. All authors reviewed the final manuscript.

## Disclosures

AS and PM have received speaker honoraria from Merck KGaA, Darmstadt, Germany. PCh has received investigator research support, consultant and speaker honoraria from Merck KGaA, Darmstadt, Germany. PCl had received research investigator support and speaker honoraria from Merck KGaA, Darmstadt, Germany. EK is an employee of Merck KGaA, Darmstadt, Germany.

### Data

All transcriptomic data will be available from Gene Expression Omnibus (GEO) and is currently in submission. Most of the GHD patient transcriptomic data is already available from GEO - GSE72439.

